# SpotMAX: a generalist framework for multi-dimensional automatic spot detection and quantification

**DOI:** 10.1101/2024.10.22.619610

**Authors:** Francesco Padovani, Ivana Čavka, Ana Rita Rodrigues Neves, Cristina Piñeiro López, Nada Al-Refaie, Leonardo Bolcato, Dimitra Chatzitheodoridou, Yagya Chadha, Xiaofeng A. Su, Jette Lengefeld, Daphne S. Cabianca, Simone Köhler, Kurt M. Schmoller

## Abstract

The analysis of spot-like structures is a widespread task in microscopy-based cell biology. Existing solutions are typically specific to single applications and do not use multi-dimensional information from 5D datasets. Therefore, experimental scientists often resort to subjective manual annotation. Here, we present SpotMAX, a generalist AI-driven framework for automated spot detection and quantification. SpotMAX leverages the full scope of multi-dimensional datasets with an easy-to-use interface and an embedded framework for cell segmentation and tracking. SpotMAX outperforms state-of-the-art tools, and in some cases, even expert human annotators. We applied SpotMAX across diverse experimental questions, ranging from meiotic crossover events in *C. elegans* to mitochondrial DNA dynamics in *S. cerevisiae* and telomere length in mouse stem cells, leading to new biological insights. With its flexibility in integrating AI workflows, we anticipate that SpotMAX will become the standard for spot analysis in microscopy data.

Source code: https://github.com/SchmollerLab/SpotMAX

## Introduction

Spot-like structures are a ubiquitous feature in fluorescence microscopy data and their analysis is a fundamental task in bioimage analysis. However, no generalist gold-standard tool for spot detection and quantification exists. Bioimage analysis has seen a revolution with the advent of AI-based workflows, but spot detection and quantification have not fully taken advantage of this potential. Current state-of-the-art (SOTA) tools are mainly developed for detecting diffraction-limited RNA transcripts from single-molecule FISH data or similar approaches, including imaging-based spatial transcriptomics^1–8^. These tools are often limited to spot detection, leaving the quantification of spot features, such as their size and intensities, as a downstream process to be implemented separately. Furthermore, microscopy data is often multi-dimensional, where the spot analysis must be performed in the multi-channel context or requires 3D detection and the ability to go beyond single time point detection. Some of the most recent SOTA tools implement advanced AI-based workflows^2,6,7^, but they remain limited to detection in 2D and none of the existing tools covers all aspects required for the rigorous analysis of spots in 5D microscopy data. This makes it hard for experimentalists to find an appropriate tool because integrating the multi-dimensional information requires advanced technical knowledge. In many cases, manual detection remains the only viable method for experimental biologists, a strategy that can be extremely tedious and time-consuming potentially introducing mistakes due to human bias. Therefore, a standard tool for generalist spot detection and quantification of 3D globular-like structures with support for time-lapse data is urgently needed.

To solve these issues, we developed SpotMAX (Spot detection by local MAX search), a generalist spot detection and quantification framework for multi-dimensional microscopy data. SpotMAX provides an easy-to-use unified framework to access AI-driven spot detection workflows. It harnesses all the relevant information available in multi-dimensional datasets, such as single-cell segmentation masks, reference channels (e.g., separate staining of the cell nucleus), and multi-dimensionality such as z-slices and time-lapse. We developed SpotMAX with three main objectives: 1) providing better performance than existing SOTA tools, 2) enabling experimental biologists to uncover new biological insights, and 3) making it easy to use without specialist programming knowledge. First, to assess the accuracy of spot detection with SpotMAX, we manually annotated a large 3D dataset of experimental data and, since such a resource was missing, we make the dataset available to the community. Next, we extensively benchmarked SpotMAX, and found that it outperforms SOTA tools, non-expert annotators, and, in some cases, even expert annotators.

Moreover, when spot counting alone is not sufficient, further quantification and information from additional channels are required. To achieve this, SpotMAX provides a framework for segmenting the reference structures and quantifies several spot features, such as their size, intensities, and signal-to-noise ratio. To test SpotMAX in real-world scenarios, we applied SpotMAX to multiple image modalities, model organisms, biological questions, and experimental protocols. This included the quantification of crossover events in *C. elegans* meiosis, multi-generational dynamics of mitochondrial DNA adaptation in response to a nutrient shift in *S. cerevisiae*, telomere length as a function of cell size in hematopoietic stem cells, and Raptor depletion effects on the nucleus and nucleolus size in *C. elegans*. For each application, we demonstrated that by leveraging multi-dimensionality and extracting biologically meaningful features, SpotMAX can be used to reveal new biological insights.

SpotMAX is available to the community as open-source software and a Python package with extensive documentation (see links below). It can be executed from Python APIs or the command line and easily scales for high-throughput analysis in high-performance computing environments. Crucially, we provide an intuitive and user-friendly GUI integrated into the Cell-ACDC software^9^. Importantly, when suitable, analysis steps can be performed with any of the models available on the BioImage.IO model zoo^10^, streamlining integration with existing strategies.

Thanks to its superior performance, ease of use, and high accessibility, we envision SpotMAX to become a standard for general fluorescence spot detection and quantification tasks in biological research.

Documentation: https://spotmax.readthedocs.io/en/latest/index.html

Source code: https://github.com/SchmollerLab/SpotMAX

## Results

### Software architecture

Microscopy data with spot-like features is often multi-dimensional including multiple channels that typically represent cells or entities containing spot-like structures. In this case, the starting point for SpotMAX analysis is the segmented masks of the single cells (or other entities). These masks are generated outside of SpotMAX and can be obtained with any segmentation software such as our previously published software Cell-ACDC^9^.

SpotMAX is composed of the following modules: 1) optional semantic segmentation of the reference channel, 2) semantic segmentation of the spot channel, 3) spot detection, and 4) spot quantification (**Fig. 1**). These modules run sequentially, and, if needed, the input of each module can be replaced by the analysis with an external software. This modularity facilitates a high degree of integration with existing methods.

**Fig. 1.**
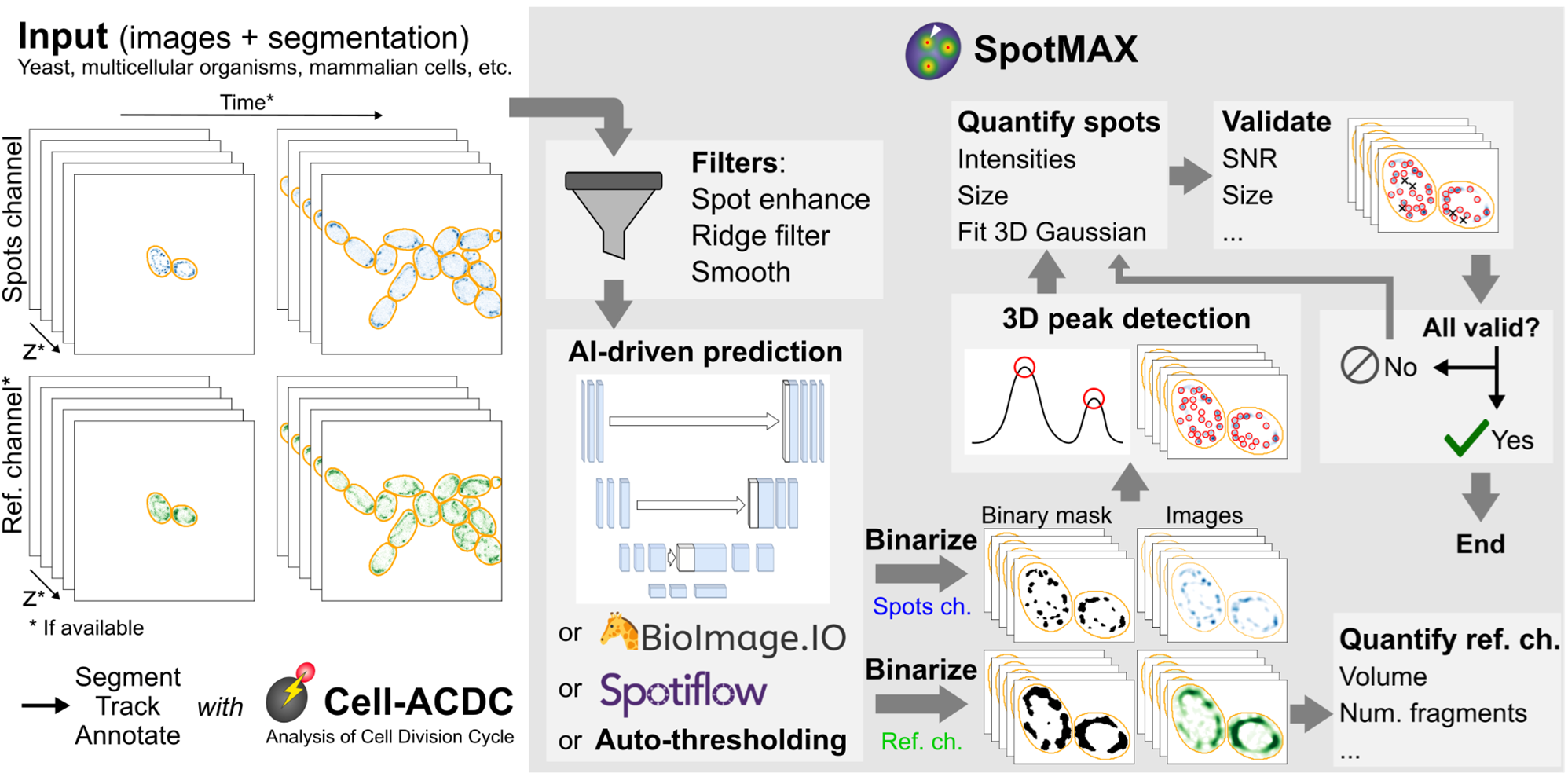
Schematic overview of the SpotMAX framework. SpotMAX leverages the multi-dimensionality of microscopy datasets, including time, z-slices, a reference channel, and segmentation masks (e.g., single cells, orange outlines). The single cells can be segmented, tracked, and mother-daughter relationships can be annotated with Cell-ACDC^9^ or any other external software. The input images plus segmentation masks go through a series of optional filters designed to enhance the structure of interest (e.g., spots for the spot channel or sub-cellular organelles for the reference channel). The pre-processed images are then fed to the prediction step, where AI-based models or automatic thresholding can be used to generate binary masks for both the spots and reference channels. These models include custom-designed 2D or 3D U-Net models (see Methods), any of the models available at the BioImage.IO model zoo^10^, Spotiflow^2^, or automatic thresholding. For the reference channel, the input segmentation masks are used to assign the binary masks to each single cell to obtain the final segmentation labels for the reference channel. These are then used in conjunction with the intensity signal to compute several features such as the reference channel structure volume, number of fragments, intensity-based features etc. The binary masks of the spot channel are used to guide a 3D peak detection pipeline that detects peaks in the signal within the masks. The detected spots are then quantified, and user-selected features can be used to filter valid spots. When false positive spots are removed, they become part of the background, hence requiring re-calculation of features such as the signal-to-noise ratio. To account for these changes, quantification and filtering are repeated until no more false positive spots are discarded.

The reference channel segmentation (module 1) is performed in two steps: a) image pre-processing, and b) semantic segmentation. For image preprocessing, two filters can be applied: a Gaussian smoothing filter and a filter that enhances network-like structures^11^. For the semantic segmentation, either automatic thresholding or any of the models available at the BioImage.IO^10^ model zoo can be applied.

For the semantic segmentation of the spot channel (module 2), the z-stack image is pre-processed with a spot detection filter (Difference of Gaussians), whose parameters are determined from the expected spot size (see Methods section). The filtered image is then used as input into 4 possible strategies: a) automatic thresholding, b) SpotMAX AI, c) Spotiflow^2^, or d) any model of the BioImage.IO model zoo. The SpotMAX AI consists of 2D or 3D U-Net-based neural networks (Supplementary Fig. S1) that we trained to perform semantic segmentation of the spot image. The training data consisted of manually annotated 3D diffraction-limited spots from single-molecule FISH^12,13^, mitochondrial DNA nucleoid data^14^, and publicly available 2D datasets^6^.

To separate the spots merged in the semantic segmentation mask and detect their centre (module 3), SpotMAX performs local peak detection with a footprint determined from the expected spot size. For diffraction-limited spots, the expected spot size is calculated automatically from the image acquisition parameters (see Methods). For larger spots, the spot size can be visually adjusted in the GUI.

To quantify spot features (module 4), SpotMAX uses the pixels of a spheroid mask with an expected spot size centred at the spot centre. These features include several intensity distribution metrics (mean, max, median, quantiles, etc.) from the raw, Gaussian-filtered, and DoG-filtered images including different background correction strategies.

Spots can also be further quantified with a Gaussian peak fitting procedure (SpotFIT, module 4). We optimised this procedure for high-density spots where multiple spots are fitted together with a sum of 3D Gaussian functions (see Methods). The SpotFIT framework computes additional features, including the total integral of the Gaussian function, the size of the spot, the amplitude, and the background level.

### SpotMAX outperforms SOTA models and human annotators

To evaluate the performance of SpotMAX in experimental datasets, we designed several benchmarks with realistic simulations of the typical annotation process faced by experimental biologists (**Fig. 2**). Because a high-quality 3D ground truth dataset did not exist so far, we asked expert annotators to annotate the spot centres in the following 3D datasets: 1) single-molecule RNA FISH in *S. cerevisiae,* 2) mitochondrial DNA (mtDNA) nucleoids in *S. cerevisiae*, and 3) chromosomal crossovers in the multi-cellular organism *C. elegans.* All annotations were performed using Cell-ACDC^9^. We generated a ground truth dataset of 369 volumes with 20692 annotated spots from 2894 cells, including 1060 cells without true positives (models were allowed to detect false positives on these cells). Each category has a different pixel size, experimental protocol, and imaging method. Details about the ground truth datasets are available in the methods and Supplementary Table 1. Next, we identified the best-performing strategy in each category to benchmark against SpotMAX. The identified tools are 1) RS-FISH^1^, 2) Spotiflow^2^, and 3) expert manual counting, respectively. For a fair comparison, with SpotMAX we did not use the trained U-Net but the automatic thresholding option because part of the benchmarking dataset was used for training the U-Net models (see Methods). For the first category containing smFISH data, we also asked a junior scientist with no previous smFISH experience (“Human” in **Fig. 2c**) to annotate the centre of the spots. Interestingly, we found that SpotMAX outperformed not only the SOTA tool but also the non-expert human annotator (**Fig. 2b**). In the second category, the spots are mtDNA-protein complexes (nucleoids) that contain about 1-3 copies of mtDNA^14,15^ and appear in the images as globular-like structures distributed along the mitochondrial network. For this category, we could not find a suitable tool for 3D spot detection, hence we decided to use pre-trained Spotiflow (general model), which was previously shown to outperform other SOTA tools^2^. Since Spotiflow can detect spots only in 2D images, we first compared it to SpotMAX 2D and found that SpotMAX 2D had a higher recall. By leveraging the 3D context, SpotMAX further improved both precision and recall, showing the importance of the multi-dimensional information (**Fig. 2d**). In the third category, the ground truth for the majority of objects is known *a priori.* Homologous chromosomes must form crossovers during meiosis to increase the genetic diversity of the gametes. This crossover formation is tightly regulated, and in *C. elegans*, each of the six pairs of homologous chromosomes receives exactly one crossover^16^. Therefore, we can observe six foci of the crossover marker COSA-1 in each nucleus after crossover designation^17^. Importantly, the annotator representing the SOTA strategy was not informed to expect 6 spots per cell, avoiding any bias in counting. Counting spots in 3D is tedious, and spots are easily counted twice across neighbouring z-slices. To minimize the occurrence of this type of error, we equipped Cell-ACDC with visual help where a spot is annotated on multiple neighbouring z-slices.

**Fig. 2.**
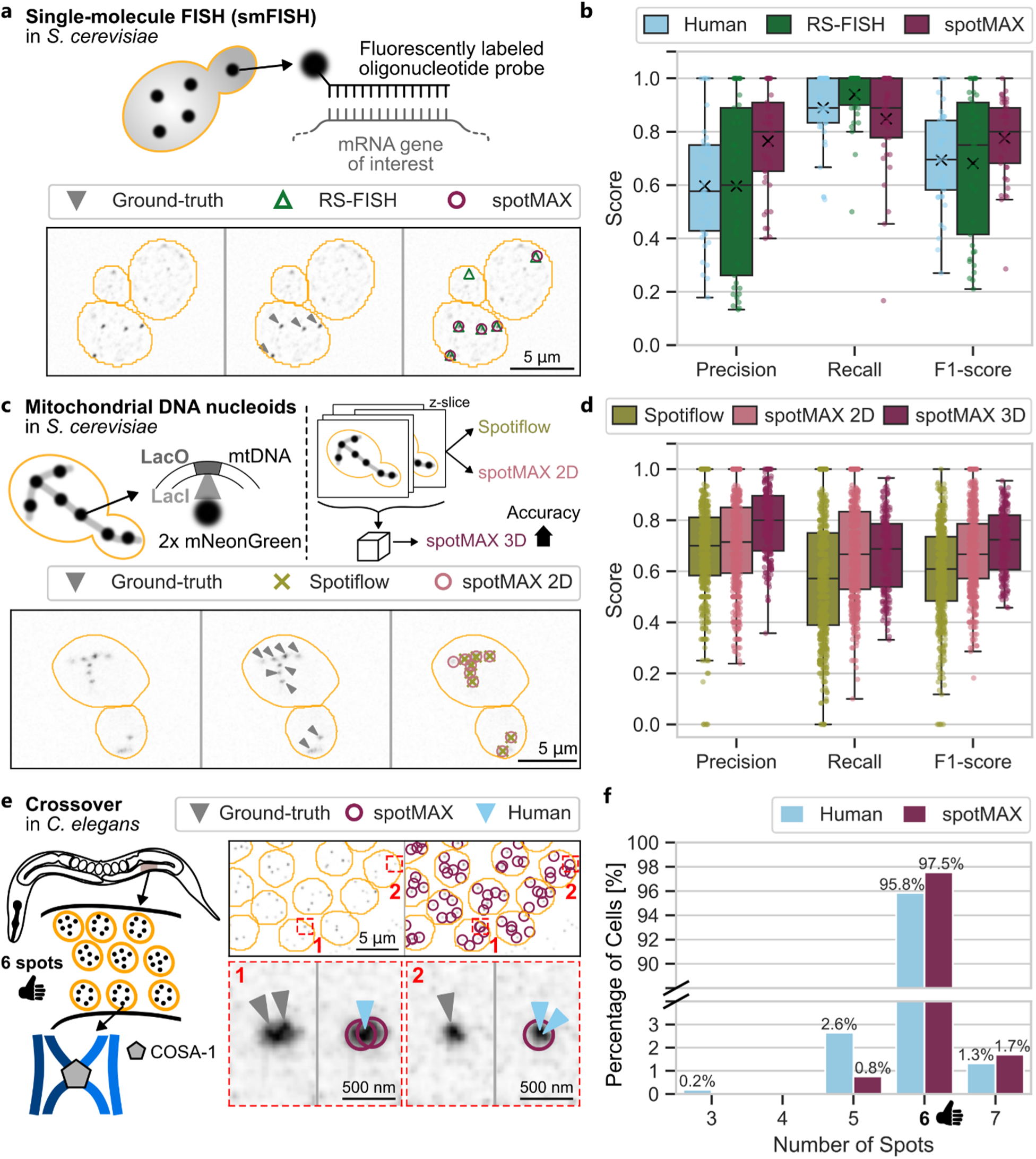
Benchmark of SpotMAX in various experimental conditions. **a**, Schematic representation of the tested dataset, single-molecule fluorescence *in situ* hybridisation (smFISH) in the model organism *Saccharomyces cerevisiae*, and representative images with ground truth, Radial Symmetry-FISH (RS-FISH) software^1^, and SpotMAX annotations. The orange outlines are the contours of the segmentation masks generated with Cell-ACDC. RS-FISH was chosen as the SOTA model in the analysis of smFISH data because it outperformed all other candidates (see ref. ^1,2^ and data not shown). **b**, Results of the benchmark between RS-FISH, Human, and SpotMAX. The ground truth data was generated by an expert in smFISH analysis in budding yeast. The “Human” annotations are from a junior scientist without previous experience with smFISH data who was instructed by the expert about the biological priors and how to annotate the data. **c**, Schematics of the tested dataset, mitochondrial DNA nucleoids counting in *S. cerevisiae* (top-left), schematics of the benchmark strategy (top-right), and representative images with ground truth, Spotiflow software, and SpotMAX 2D annotations. The orange outlines are the contours of the cell segmentation masks generated with Cell-ACDC. **d**, Results of the benchmark between SpotMAX 2D (selected z-slices as input), Spotiflow, and SpotMAX 3D (entire z-stack as input). The ground truth was annotated by an expert annotator in the field of mitochondria biology. **e**, Schematics of the tested dataset, chromosomal crossovers counting in the germline of the multi-cellular organism *Caenorhabditis elegans*, and representative images with ground truth, SpotMAX and Human annotations. The orange outlines are the contours of the segmentation masks generated with Cell-ACDC. To visualise crossovers, we engineered two new strains with either mNeonGreen or HaloTag fused to COSA-1, a key protein required for crossover formation. The ground truth in this dataset is known *a priori* because there are 6 chromosomal crossovers in almost every cell. The “Human” is the smFISH expert annotator, who had no experience with counting crossovers in *C. elegans* and did not know the expected number of spots per cell. The representative images show two instances where the human made a counting mistake. **f**, Results of the benchmark between SpotMAX and Human. As expected, both Human and SpotMAX counted 6 spots in most of the cells. However, SpotMAX had a better performance. This high level of accuracy is key to studying crossing over in *C. elegans*, as errors in crossover regulation are rare, and even a single missing or extra crossover can result in aneuploidies in the gametes. Details about the datasets used in this benchmark are available in Supplementary Table 1. Image intensities were adjusted for visualisation purposes.

When counting chromosomal crossovers in *C. elegans* (**Fig. 2e**), we found that the human annotator counted 5, and 7 spots in 2.4%, and 1.3% of the cells, respectively, compared to 1.5%, and 1.7% of SpotMAX (**Fig. 2f**). The human annotator also counted 3 spots in 0.4% of the cells (human average count = 5.98 ± 0.24 spots, SpotMAX average count = 6.01 ± 0.16 spots, where the range is the standard deviation). Surprisingly, upon closer inspection, we found that all SpotMAX counts appear correct (consistent with rare cell-to-cell variation in WT^18^). Overall, the process of crossover formation is tightly regulated, and small perturbations can have drastic effects. Any extra or missing crossovers can result in the mis-segregation of chromosomes during the meiotic divisions and produce aneuploid gametes that give rise to infertility or congenital conditions^19^. Therefore, an unbiased high-accuracy and high-throughput method was desperately needed to measure these small perturbations.

### Apoptosis targets meiotic nuclei with crossover errors, bypassing the DNA damage checkpoint

Automated and unbiased analysis is often key to answering open biological questions. For example, to study meiosis, understanding chromosomal crossover (CO) formation is fundamental because errors in this process are a major cause of infertility and can result in congenital conditions such as Down syndrome^20^. A widely used model organism in this field is *C. elegans*. The majority of the cells in the germline of this animal form 6 COs. In rare cases, very few cells will have an abnormal number of COs (**Fig. 3a,b**). To ensure the genetic integrity of the offspring, such nuclei should be promptly removed by apoptosis. However, exactly how this process works and how apoptosis is triggered under such circumstances is unknown. To investigate the rare occurrence of an aberrant number of COs in *C. elegans*, manual counting, while widely used and effective due to its low error rate, lacks the throughput necessary to study these infrequent events.

**Fig. 3.**
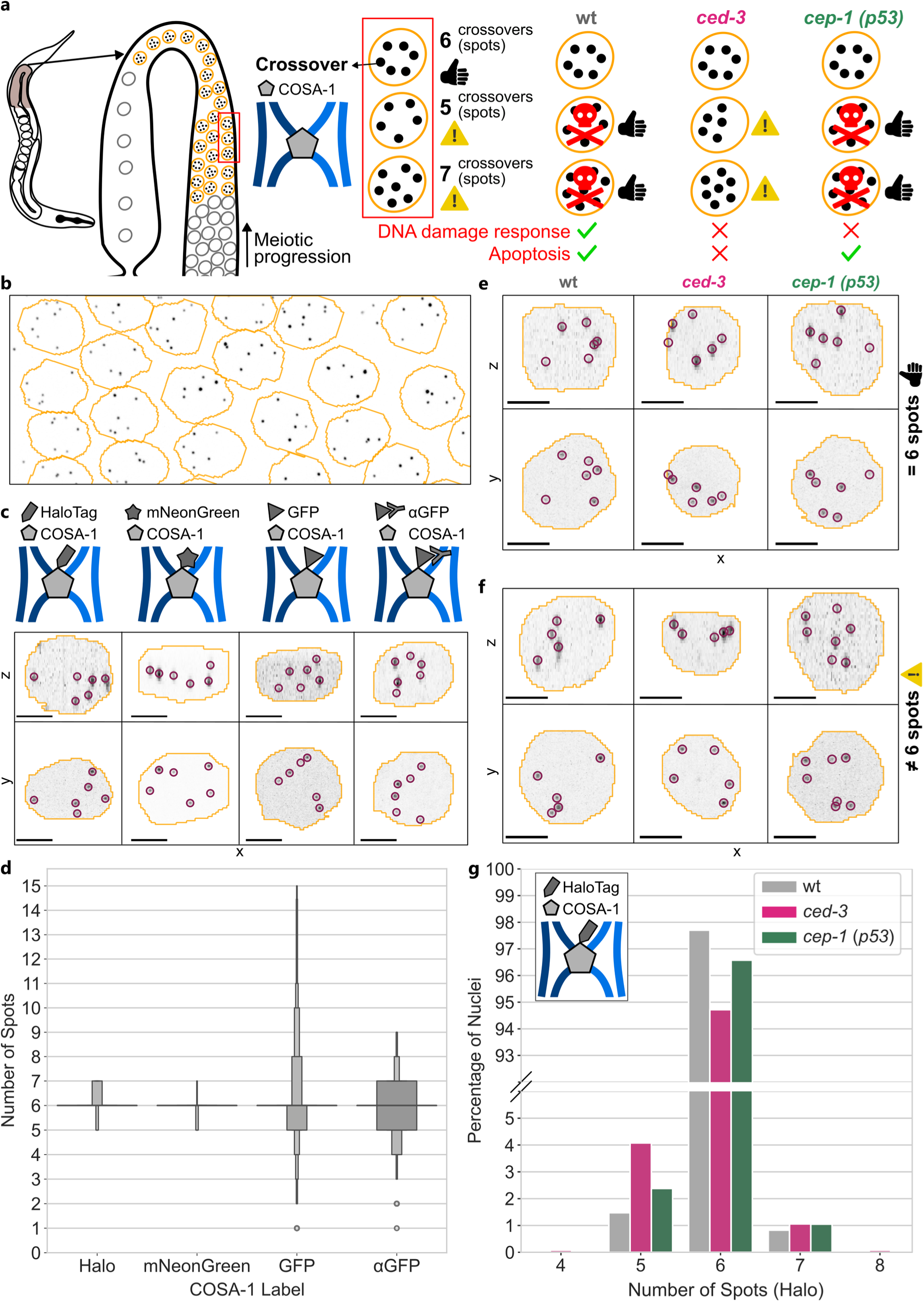
SpotMAX analysis of the crossover error regulation process in *C. elegans*. **a**, Schematic representation of the experimental setup and conditions in mutant strains. To study chromosomal crossovers (COs) in the *C. elegans* germline, we engineered new strains by either fusing mNeonGreen or a HaloTag to COSA-1, a key protein in crossover formation. The expected number of COs in wild-type cells is 6. In a small number of cells, 5 or 7 COs might form. These aberrant nuclei may be removed through apoptosis. By deleting *ced-3*, apoptosis is inhibited, resulting in more cells with an abnormal number of COs. By deleting *cep-1 (p53)*, only DNA-damage response triggered apoptosis is inhibited, but the cells with abnormal numbers of COs are still removed, indicating that other apoptosis pathways are involved in their removal. **b**, Representative image of cells within the germline from the COSA-1::mNeonGreen strain (see c panel). **c**, Schematics representation of the four methods we tested to visualise chromosomal crossovers, and representative images for each method with SpotMAX annotations (purple circles) and segmentation outlines generated with Cell-ACDC^26^. GFP, HaloTag, and mNeonGreen are fluorescent proteins fused to COSA-1, while αGFP refers to antibody staining of the GFP (gold-standard). **d**, Letter-value plots^27^ showing the number of COSA-1 spots (Number of Spots) quantified with SpotMAX for the four visualisation methods (see panel c). Number of nuclei: Halo = 453, mNeonGreen = 603, GFP = 630, αGFP = 630. **e**, Representative images of wild-type (wt), *ced-3*, and *cep-1 (p53)* deletion mutants where the number of COs is the expected value of 6. Purple circles are the SpotMAX detections, while the orange outlines are the segmentation masks generated with Cell-ACDC. **f**, Same as in panel e, but for cells that have more or less than 6 spots. **g**, Histograms of the number of COs in cells without unrepaired DNA damage (Number of Spots) counted with SpotMAX in wt, *ced-3*, and *cep-1* mutants (Halo::COSA-1 staining). Number of nuclei: wt = 1698, *ced-3* = 1325, *cep-1* = 1430. Scale bar is 2 μm. Image intensities were adjusted for visualisation purposes.

Therefore, we tested SpotMAX’s ability to accurately count spots in a large number of cells in a fully automatic and unbiased way. To visualise the COs, we first generated two *C. elegans* strains carrying the endogenous reporters HaloTag and mNeonGreen fused to the COSA-1 gene, a key component of CO formation^17^. More details about the dataset are available in Supplementary Table 2.

Next, we used SpotMAX to compare the robustness of different markers for COSA-1 by quantifying the native fluorescence of 3 reporters (GFP, mNeonGreen, and HaloTag, (**Fig. 3c**) and antibody staining of the GFP reporter. The GFP::COSA-1 reporter is expressed from a transgene generated by microparticle bombardment. Halo::COSA-1 and COSA-1::mNeonGreen are endogenously tagged by CRISPR/Cas9 mediated genome editing (see Methods) and are fully functional (Supplementary Table 4). mNeonGreen surpasses the commonly used GFP::COSA-1 reporter in brightness and signal-to-noise ratio, while the spot intensities of Halo::COSA-1 are the least variable (Supplementary Fig. S2a), which directly translates to better accuracy in the SpotMAX quantification (**Fig. 3d**). By contrast, antibody staining of GFP::COSA-1 introduces a surplus of nuclei with more than 6 foci due to an increase in background staining resulting in a lower signal-to-noise ratio.

The fact that 97% of meiotic cells designate exactly one crossover on each of the six pairs of homologs, resulting in six spots per nucleus, highlights the robustness of the regulatory mechanisms controlling crossover number and distribution during meiosis. Alternatively, or additionally, there may be a checkpoint that eliminates nuclei with an incorrect number of crossovers. Indeed, 50% of cells are removed through apoptosis during meiotic prophase in *C. elegans*^19^, and persistent defects in the meiotic program can activate apoptosis of meiotic cells across metazoans^20^. We wondered if nuclei with abnormal numbers of designated crossovers are preferentially targeted for apoptosis to enhance the robustness of crossover regulation. Therefore, we compared the percentage of nuclei with aberrant numbers of Halo::COSA-1 foci in wild-type and *ced-3* animals (**Fig. 3a,e,f**), which cannot initiate programmed cell death in both somatic and germline cells^21^. In wild-type animals, only 2.7% of nuclei had aberrant numbers of COSA-1 foci, compared to 6.5% in *ced-3* animals (Supplementary Fig. S2b). Most nuclei with aberrant COSA-1 foci had five or fewer foci, suggesting some chromosomes lacked a crossover. Crossover assurance promotes double-strand break formation until each chromosome pair has at least one crossover^22^. The absence of a crossover on some chromosomes might result from incomplete repair of breaks into crossover precursors, which should trigger apoptosis via the DNA damage checkpoint. We tested whether *cep-1 (p53)* animals, which are deficient in DNA damage-induced apoptosis^23,24^, exhibited similar aberrant COSA-1 foci numbers as *ced-3* animals. Interestingly, *cep-1 (p53)* animals had 4.1% of nuclei with aberrant COSA-1 foci, lower than *ced-3* animals but slightly higher than wild-type animals (Supplementary Fig. S2b).

This finding suggests that nuclei with aberrant numbers of designated crossovers are targeted for apoptosis both through activation of the DNA damage checkpoint and independently of it. To test this hypothesis, we quantified spot numbers selectively in nuclei that did not have any markers of residual DNA damage. Specifically, using SpotMAX, we identified nuclei with remaining unrepaired breaks^25^, visualised by staining for V5::RAD-51, and excluded these nuclei from our analysis. In this refined analysis, *cep-1 (p53)* animals displayed similar numbers of COSA-1 foci per nucleus as wild-type animals (wild-type: 97.3% nuclei with 6 spots (5.99±0.14 spots), *cep-1 (p53)*: 95.9% nuclei with 6 spots (5.99±0.18 spots); Student t-test p-value of WT-vs-*cep-1* = 0.298). However, *ced-3* animals still had significantly fewer nuclei with six COSA-1 foci (94.7% (5.97±0.24 spots), Student t-test p-value of WT-vs-*ced-3* = 0.0015) (**Fig. 3g**). Thus, the high-throughput automated analysis of thousands of nuclei with SpotMAX revealed an unexpected mechanism that selectively targets nuclei with errors in crossover regulation for apoptosis independently of the DNA damage checkpoint.

### Asymmetric inheritance promotes fast adaptation of mtDNA copy number in *S. cerevisiae* daughter cells

So far, we tested SpotMAX’s ability to use the 3D information of z-stack volumes in a single fluorescence microscopy channel. However, many microscopy datasets are 5-dimensional including multiple channels, such as bright-field or phase contrast for tracking single cells, additional fluorescence channels (e.g., organelles), and the time dimension. Therefore, we reasoned that including information from these additional dimensions and channels could improve the overall accuracy and provide new biological insights. With time-lapse microscopy data, crucial information comes from cell identity and cross-generational pedigree information. Therefore, we integrated SpotMAX into the Cell-ACDC framework that enables segmentation, tracking, and cell-pedigree annotation in a format fully usable by SpotMAX. Additionally, to test SpotMAX’s capabilities to accurately quantify sub-cellular structures in 4D microscopy time-lapse data, we asked whether it could be used to reveal the dynamic adaptation of budding yeast mitochondria to a nutrient shift. Budding yeast can adopt two different types of metabolism, respiration and fermentation, and it upregulates the copy number of its mitochondrial DNA (mtDNA) and the mitochondrial network volume when it respires^14,28^. Yeast cells perform mainly respiration requiring mtDNA when provided with galactose as the sole carbon source^29^. We thus expect a higher mtDNA copy number and mitochondrial network volume than in cells growing on glucose media, where they can rely on anaerobic fermentation independent of mtDNA. To follow the dynamics of this adaptation, we imaged live budding yeast single cells for over 7 hours going through a medium change, from respiratory medium (SCGal, 2% galactose) to fermentation medium (SCD, 2% glucose). To visualise the mitochondrial network and mtDNA, we used a previously established system based on LacO arrays stably integrated into the mitochondrial genome and endogenously expressed fluorescent reporter proteins^14,30^ (see strain FPY004 in Supplementary Table 3). This system allows us to visualise the mitochondrial network (green in **Fig. 4a,b**) and the mitochondrial DNA (magenta in **Fig. 4a,b**, spot detections in orange circles). More details about the dataset are available in Supplementary Table 2. After data acquisition, we segmented and tracked the cells, annotated the cell cycle progression, and calculated the cell volume from the bright-field channel using Cell-ACDC^9^. The tracked segmentation masks were used as input for SpotMAX, which automatically segmented the mitochondrial network, quantified its volume, and counted the number of spots in each cell at every time point. Using the cell division annotation, we followed the evolution of the mitochondrial network volume and the nucleoid number at a given cell cycle stage (**Fig. 4c,d**). We found that the concentrations of mitochondria network volume and nucleoids directly after cell division decreased in both mother and daughter cells after shifting from SCD to SCGal. However, the concentration in the daughter cells decreases faster than in the mother cells. This differential regulation in mother and daughter cells was observed at the first time point after cell division. Since other types of regulation such as mitophagy are unlikely to happen on this short time scale (15 minutes), our findings suggest that yeast achieves a rapid adaptation of daughter cells to the new nutrient condition through asymmetric inheritance of the mitochondria and mitochondrial DNA.

**Fig. 4.**
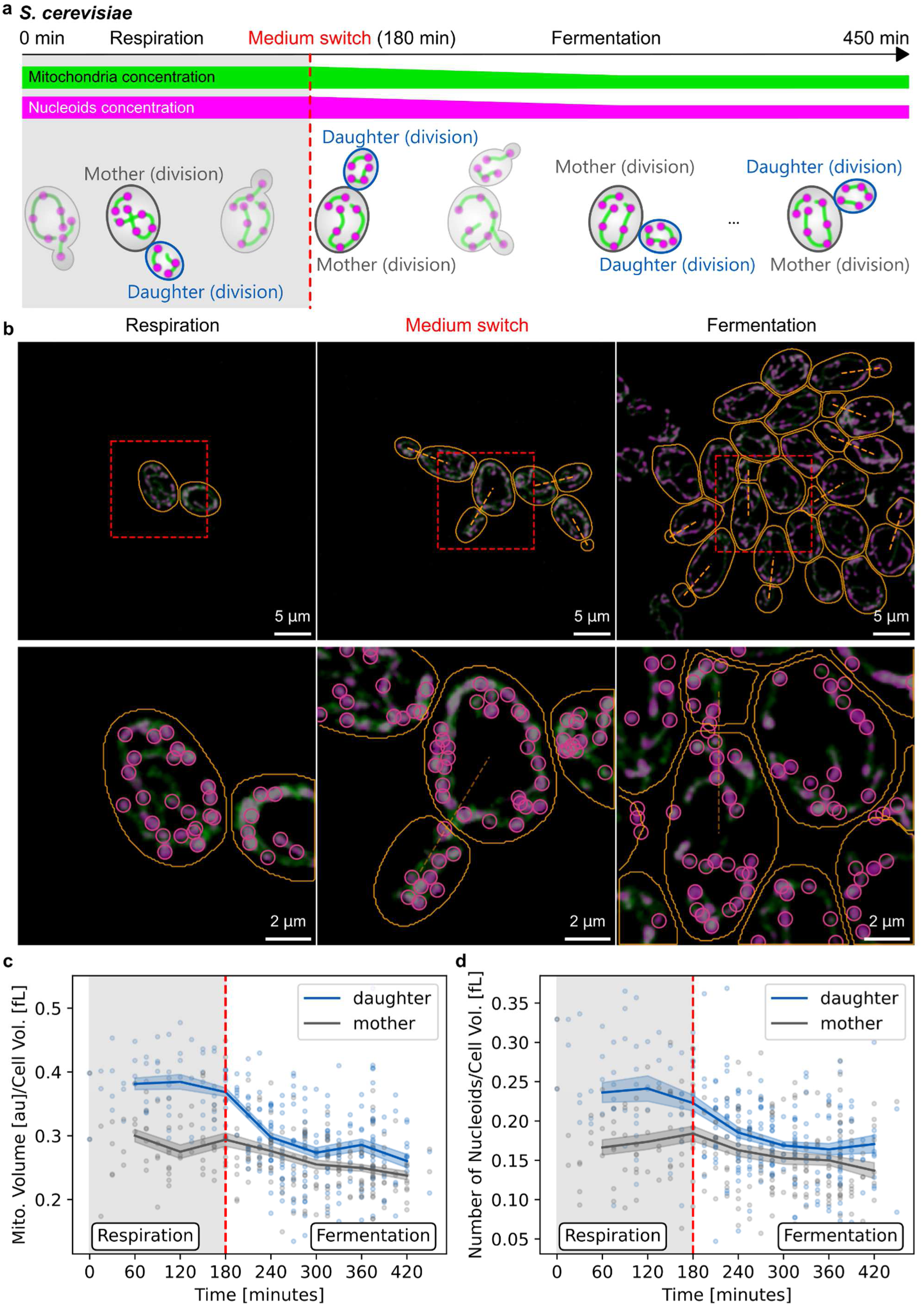
Mitochondria and nucleoid concentration in live *S. cerevisiae* cells during a nutrient shift. **a**, Schematic representation of the experimental conditions. To test SpotMAX’s capability to work with dual-channel time-lapse microscopy data, we imaged *S. cerevisiae* cells in a microfluidics chamber with an automatic medium switch. After growing cells in a medium where respiration is the preferred metabolism (2% galactose), cells were imaged for 180 minutes in the same respiration medium. After that, the medium was automatically switched to a medium containing 2% glucose, which leads to fermentation. While it is known that budding yeast cells have a higher concentration of both mitochondria and mitochondrial DNA in respiratory conditions, how this adaptation is achieved is poorly understood. After segmenting, tracking, and annotating mother-daughter relationships with Cell-ACDC, we computed the mitochondrial network volume from the reference channel and the number of nucleoids (spots), each directly after the time of division for mother cells and their daughters. We used the same visualisation approach as depicted in Fig. 2c. **b**, Representative images of budding yeast cells with mitochondrial network (green) and nucleoids (magenta) channel from three time-points: start of the experiment (respiration), medium switch, and end of the experiment (fermentation). The orange outlines show the segmentation masks, while the orange dashed line shows the mother-daughter relationships. The zoomed images show the SpotMAX annotations (pink circles) of the cells present in the first frame and their relative daughters. **c**, Mitochondrial network volume and **d**, the number of nucleoids divided by cell volume (concentration) for the cells at division as a function of time since the start of the experiment. Note that we plotted only those cells that were present at the medium switch (red dotted line). Points show data corresponding to individual cells. The line represents the mean, and the shaded area is the 95% confidence interval of the mean calculated by bootstrapping (1000 bootstrap resamples). Details about the dataset size are available in Supplementary Table 2. Image intensities were adjusted for visualisation purposes.

### Telomere length is unchanged in large hematopoietic stem cells

In many applications, spot counting is not enough, and we need to quantify the intensities of the spots. To achieve this, we equipped SpotMAX with the SpotFIT module which provides the ability to quantify the total fluorescence intensity of each spot (see Methods). To test this functionality in a biological application, we focussed on the relationship between telomere length and cell size. The declining function of hematopoietic stem cells (HSCs) during ageing is associated with an increase in their cell size^31^, but how cell size affects HSC functionality is unknown. One potential explanation is telomere shortening, which is linked to cellular ageing and causes senescence and apoptosis of somatic cells^32,33^. However, whether this applies also to HSCs is still unclear^34^. For example, during division of HSCs, telomeres become shorter^35^. Nevertheless, preventing this shortening by telomerase overexpression does not prevent HSC exhaustion in mice^34^. To further evaluate the role of telomere shortening in HSC ageing, we analysed telomere length in small, medium, and large HSCs using DNA-FISH against telomeric repeats. Next, we used SpotMAX to detect telomeres and calculated integrated spot intensities since they correlate with telomere length^36^ (**Fig. 5a,b**). As a control for which we did not expect changes in spot intensities, we stained centromeres. We used Cell-ACDC to segment HSCs and then automatically quantified telomere and centromere spot numbers and spot intensities using SpotMAX. More details about the dataset are available in Supplementary Table 2.

**Fig. 5.**
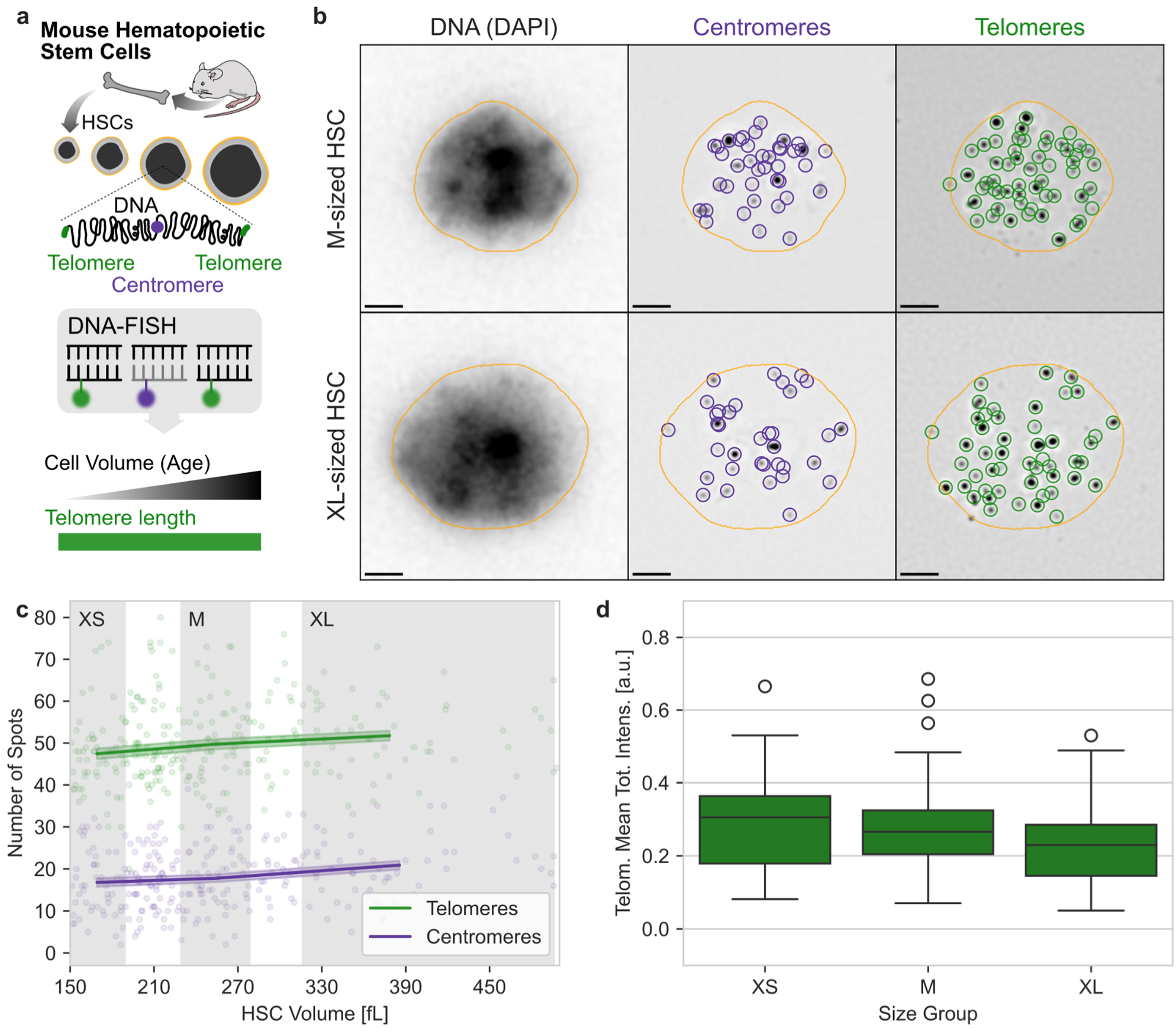
Telomeres and centromeres visualised with DNA-FISH in hematopoietic stem cells. **a**, Schematics of the experimental protocol. After extracting hematopoietic stem cells (HSCs) from mice bone marrow, we visualised the centromeres and telomeres with DNA-FISH. **b**, Representative images of medium-sized (M) and extra-large (XL) HSCs from wild-type mice showing DNA (DAPI), the telomere (Cy3, purple circles) and centromere (Alexa488, green circles) DNA-FISH signal. Circles indicate the SpotMAX detections, and the orange outlines show the cell segmentation masks obtained with Cell-ACDC^9^. **c**, Number of spots as a function of HSC volume (fL) for telomeres and centromeres. Gates of XS-, M-, and XL-sized HSCs are indicated in grey. Cells with volume less than 130 fL were discarded from the analysis because the number of spots was variable with cell size indicating a detection limit in smaller cells. The full analysis is shown in Supplementary Fig. S3a. Points in the plot represent single-cell data, the line shows the mean and the shaded area is the 95% confidence interval of the mean calculated with 1000 bootstrap resamples. **d**, Mean total intensity of the spots in the single cells for each size category. Number of cells per category: XS = 54, M = 65, XL = 54, Total = 173. The difference between the groups is not statistically significant (T-test, Bonferroni adjusted p-values: XL-vs-XS=0.186, M-vs-XS=0.697, M-vs-XL=0.186). The total intensity was calculated by integrating the Gaussian curve fitted to the data in the SpotFIT step of the SpotMAX analysis (see Methods). The total intensity of the spots in each single cell was then averaged to obtain the mean total intensity. Scale bar is 2 μm. Image intensities were adjusted for visualisation purposes.

SpotMAX detected 18 centromeric spots of variable brightness (on average, purple in **Fig. 5c**) in line with previously reported values^37^. The high variability is likely due to centromere clustering, but the number of centromeric spots stayed constant with increasing cell size (**Fig. 5c** and Supplementary Fig. S3a), suggesting that cell size does not affect their detection. Similarly, the number of detected telomere spots was unchanged in larger HSCs. Next, we investigated the mean total intensity of the telomere and centromere spots (**Fig. 5d** and Supplementary Fig. S3b). We observed that telomere length did not change in enlarged HSCs suggesting that their dysfunction is not caused by telomere shortening (**Fig. 5d**). Overall, these results demonstrate that SpotMAX enables the reliable and efficient detection and analysis of fluorescence spots of murine stem cells. Furthermore, our data suggest that telomere length is not a major contributor to the cellular ageing of enlarged HSCs.

### Nucleolus and nuclear size decrease upon DAF-15 (Raptor) depletion in the *C. elegans* intestine

Next, we asked whether SpotMAX can be used to quantify the volume of larger cellular structures in 3D microscopy data of a complex, multi-tissue organism. Specifically, we decided to test the quantification of the nucleolus and again chose *C. elegans* as a model. Nucleolar size typically reflects the activity of ribosome biogenesis^38,39^. In fact, nucleolar size is regulated by the mechanistic target of the rapamycin (mTOR) signalling pathway^40,41^. mTOR is a master regulator of ribosome production, particularly through the mTOR complex 1 (mTORC1), which regulates cell growth in response to nutrients^42^. Thus, to test the ability of SpotMAX to quantify changes in nucleolar volume, we used animals expressing the nucleolar marker FIB-1::mCherry (Fibrillarin)^40^, as well as HIS-72::GFP (H3.3) to label nuclei. We used the auxin-inducible degradation (AID) system^43^ to degrade DAF-15, the *C. elegans* homolog of mammalian Raptor, a specific component of mTORC1^44^. More details about the dataset are available in Supplementary Table 2. We imaged larvae treated with auxin or ethanol as a control, and then segmented intestinal nuclei with Cell-ACDC^9^, using the HIS-72::GFP (H3.3) signal at the central z-slice of each nucleus. These masks were then used as input for SpotMAX (which automatically expanded the 2D segmentation masks 10 z-slices above and below the segmented one) to detect and quantify the volume of the corresponding nucleoli (**Fig. 6a,b**). Consistent with previous studies that directly perturbed the LET-363 (mTOR) kinase^40,41^, we found that depletion of DAF-15 drastically decreases nucleolar volume (**Fig. 6c**).

**Fig. 6.**
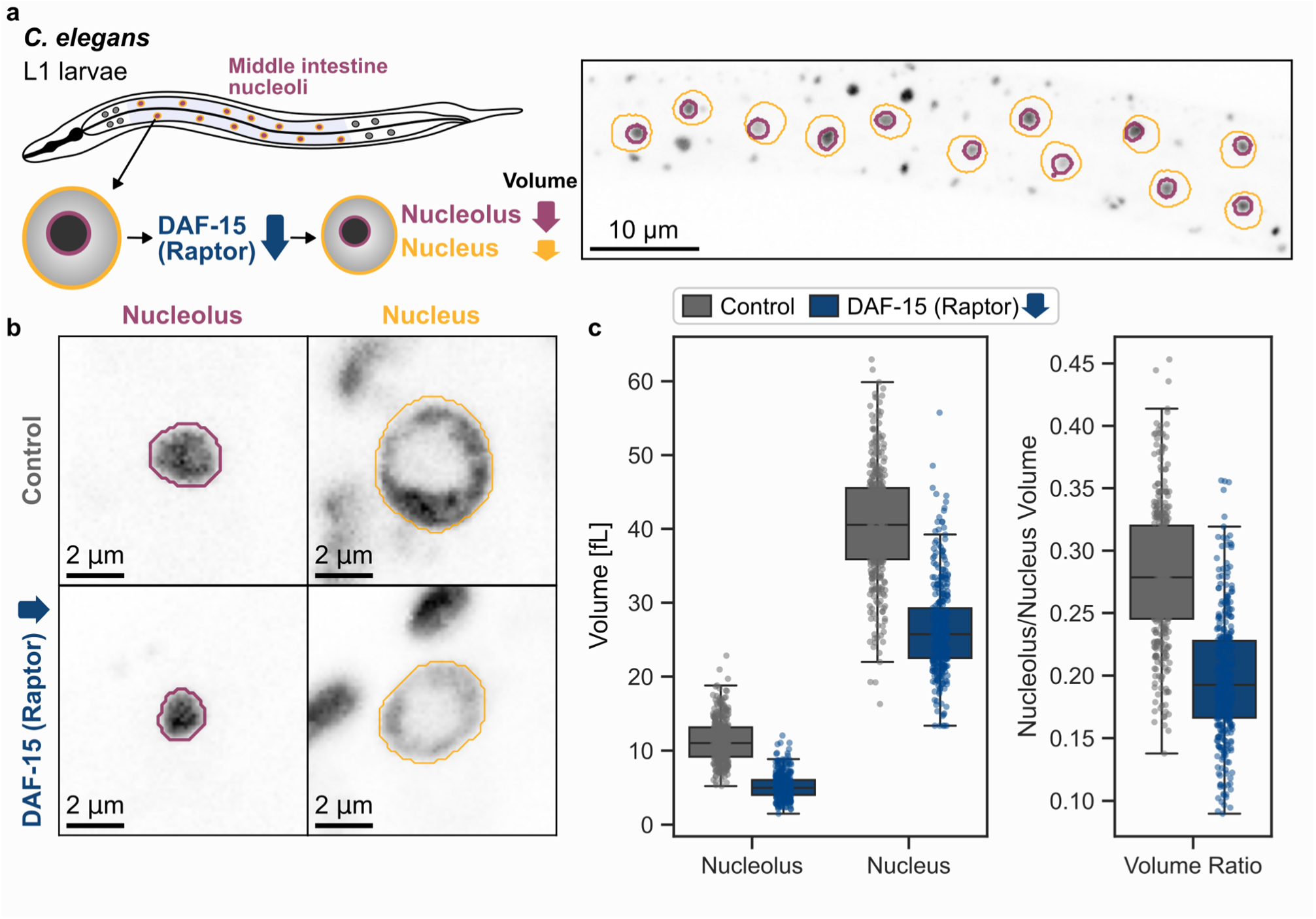
Nucleolus and nucleus size upon DAF-15 (Raptor) depletion in the *C. elegans* intestine. **a**, Schematic representation of the experimental protocol. To test SpotMAX’s capabilities to segment globular structures and compute their volume, we quantified the nucleolus and nucleus volume decrease upon DAF-15 (Raptor) depletion in the 12 middle intestine cells in *C. elegans* L1 larvae. The representative image on the left shows the 12 intestinal nuclei segmented with Cell-ACDC (orange outlines) in the nucleoli channel. Purple outlines represent SpotMAX’s automatic segmentation of the nucleoli (spots). **b**, Representative images of the nucleolus (purple outlines, calculated by SpotMAX) and the nucleus (orange outlines from Cell-ACDC) channels in both the control and DAF-15 depleted strains. **c**, Volume of the nucleolus, the nucleus and the nucleolus/nucleus ratio in the control and DAF-15 depleted strains. Student’s t-test p-values WT-vs-DAF-15 for Nucleolus, Nucleus, and Volume Ratio are all <10^-10^. Details about the dataset size are available in Supplementary Table 2. Image intensities were adjusted for visualisation purposes.

While the impact of mTOR on nucleolar size is well documented in different biological systems^38,40,45^, it is less clear whether it also affects nuclear size. Using the segmentation masks described above, we quantified the HIS-72/H3.3-GFP signal as a readout for nuclear size and found that the volume of intestinal nuclei is also reduced upon DAF-15 degradation (**Fig. 6c**). However, the relative reduction in size is greater for the nucleolus compared to the nucleus, revealing that intestinal cells of mTORC1-inhibited animals have a reduced nucleolus-to-nucleus volume ratio.

## Discussion

Here, we presented SpotMAX, a generalist 3D spot detection and quantification tool for the analysis of fluorescence microscopy data. SpotMAX introduces three main innovations: 1) it is versatile and application-independent, 2) it can leverage multi-dimensional information, including multiple fluorescence channels, z-stacks, and time information, and 3) it goes beyond spot detection by computing biologically meaningful features. Along with SpotMAX, we also publish a high-quality 3D ground truth dataset of experimental data from various experimental protocols and imaging modalities composed of 369 z-stack volumes with 20692 annotated spots from 2894 cells. This dataset is the first public dataset of its kind, and it can therefore serve as a reference to test future spot detection tools. We showed that SpotMAX outperforms current state-of-the-art tools and can provide new biological insights, a result made possible by several improvements compared to existing tools. First, SpotMAX employs both AI-based and traditional computer vision algorithms, including a custom-made 3D U-Net architecture and existing AI tools such as Spotiflow and any of the models available on the BioImage.IO zoo^10^. Paired with a user-friendly GUI and tight integration with Cell-ACDC^9^ as well as other existing tools, we provide a framework for experimental biologists to tackle open biological questions without advanced technical knowledge of complex bioimage analysis pipelines. Secondly, SpotMAX goes beyond spot detection by computing several morphological and intensity-based features that allowed us to achieve a significant performance increase. This drastically simplifies the building of image analysis pipelines where spot counting is not enough.

Finally, to meet the current demands of running workflows on high-performance computing (HPC) environments, we provide both the possibility to run SpotMAX headless from the command line as well as from Python APIs and provide extensive documentation. This can be especially valuable to software developers and bioimage analysts who need to integrate spot detection and quantification algorithms within existing workflows. Furthermore, the analysis of multiple images can be readily parallelised in HPC environments.

We believe that SpotMAX’s ability to go beyond state-of-the-art spot detection with a framework for accurate multi-channel and multi-dimensional quantification and its tight integration with current AI pipelines will make it a reference software for experimental biologists addressing spot quantification tasks.

## Methods

SpotMAX is written in Python v3 and published on the Python Package Index repository. We extensively documented SpotMAX, including two tutorials with experimental data. The documentation can be found at https://spotmax.readthedocs.io/en/latest/index.html and the source code is hosted on GitHub at https://github.com/SchmollerLab/SpotMAX. SpotMAX can be used as a plugin of Cell-ACDC^9^ through a user-friendly GUI written with QtPy and PyQtGraph^40^. Alternatively, it can be deployed in a high-performance computing environment from the command line or with the Python APIs.

### SpotMAX AI architecture and training

The SpotMAX AI model comprises 2D and 3D models with a U-Net-style architecture written with PyTorch^41^. The 2D model is custom-made, while the 3D model is based on the library pytorch-3dunet^42^.

#### Models

The 2D and 3D model architectures are shown in Supplementary Fig. S1. Both models employ an encoder-decoder architecture with a sequence of batch normalization, convolution layers, ReLU and max pooling. Additionally, each feature map of the encoder is concatenated to its corresponding feature map of the decoder. The final step consists of a convolution layer with a 1x1 kernel size to obtain the two classes, the background and the foreground. A pixel in the foreground is a pixel predicted as belonging to a spot (i.e. semantic segmentation).

#### Datasets and ground truth generation

The ground truth data for the training of the SpotMAX AI models consists of binary masks of the spot signal of various datasets. Specifically, we used previously published 3D datasets analysed with Cell-ACDC^12–14^ (for both 3D and 2D models), as well as the 2D “smfish” and “suntag” datasets used in the deepBlink software (for the 2D model)^6^. To generate the spot masks, we started from the (x, y, z) spot coordinates provided by the corresponding publications and we generated binary masks by placing a spheroid with fixed radii (different z-radius to accommodate anisotropic volumes) in each spot centre. The radii were calculated using the Abbe diffraction limit formula in x and y and multiplied by 3 to obtain the radius in z. These masks were manually inspected and, if necessary, corrected by an independent expert annotator using the Cell-ACDC software. The dataset was then split into train, validation, and test sets. The final dataset is composed of 4956 training z-slices, 1471 validation z-slices, and 1586 test z-slices.

#### Image preprocessing

Before feeding the model, the images are pre-processed in three steps: 1) resizing based on the pixel size to match a final pixel size of 73 nm/pixel; 2) grayscale morphological opening to remove hot pixels artefacts (isolated bright pixels); 3) MinMax scaler, which scales each image to the (-1, 1) range based on the global minimum and maximum in each dataset.

#### Training

Both models were trained using stochastic gradient descent. The 2D model was trained for 100 epochs with a batch size of 16 images cropped to 400x400 size. Since the information in the z-direction is not encoded by the model, z-slices were randomly shuffled before entering the model. For the loss function, we used the inverse of the Dice score. For the learning rate, we used a scheduler that, starting from a value of 0.001, reduces the rate after the validation score stops improving for more than eight epochs. As a result, the learning rate started decreasing around the 25 epochs and reached a final value of 1e-8.

For the 3D model, we divided the datasets into patches, to reduce the computational cost, which can be high when working with 3D convolutional layers. The patches were extracted using a (z, y, x) patch size of (30, 250, 250) pixels with a stride of (20, 100, 100). Note that we tested several patch and stride sizes (data not shown) and selected the ones that resulted in the best performance. Furthermore, unlike with the 2D model, z-slices were not shuffled, since the correct order is important for the 3D operations. Finally, we used a batch size of 4 and 15000 training steps with the same learning rate scheduler used for training the 2D model.

All training runs were performed on a GPU Nvidia Tesla V100 with 32 GB on the BMBF-funded de.NBI Cloud within the German Network for Bioinformatics Infrastructure. The training time for a single run was around 5 hours for the 2D model and 8 hours for the 3D.

### 3D Gaussian fitting procedure (SpotFIT)

To quantify spots with an intensity profile similar to a normal distribution, SpotMAX fits 3D Gaussian functions to the spots’ signal. The 3D Gaussian function is defined as follows:

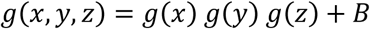

where *B* is a fitting parameter and *g*(*x*), *g*(*y*), and *g*(*z*) are 1D Gaussian functions given by

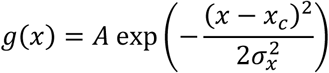

where *x_c_*, *σ_x_*, and *A*, are fitting parameters and they are the spot centre coordinate, the width (sigma), and the amplitude of the Gaussian peak. The integrated spot intensity used in the telomere length analysis in hematopoietic stem cells is calculated as follows:

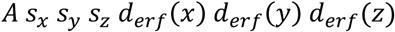

where 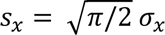 and the same for the other dimensions, and *d_erf_*(*x*) is defined as follows:

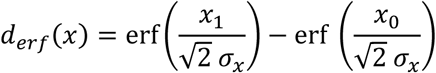

with erf being the error function. *x*_1_ and *x*_0_ are the upper and lower bounds coordinates around the peak centre and they are *x*_1_ = 1.96 *σ_x_* and *x*_0_ = −1.96 *σ_x_*. The same is then calculated for the *y* and the *z* directions.

Since fitting all pixels in the image is computationally infeasible and spots are often close to each other, we designed a workflow to determine relevant pixels. To account for differences in spot sizes, SpotMAX identifies the relevant pixels for each spot by iteratively enlarging the spheroid footprints by one pixel in each direction until the mean of the outer pixels in the spheroid mask falls below a threshold. This threshold is calculated as the background median plus three times the background standard deviation. Next, during the SpotFIT routine, SpotMAX fits one or more 3D Gaussian peaks to the spot intensities in the previously determined spot masks. To achieve the highest accuracy possible, with N touching spots, the sum of N Gaussian peaks should be fitted. However, this becomes extremely computationally intensive with an increasing number of touching spots. Therefore, we implemented a computationally efficient procedure where the number of Gaussian peaks fitted is minimized. Given a group of touching spots, SpotMAX sorts the spots based on the number of direct neighbours in ascending order and starts from the spot that touches the least number of spots and runs the fitting procedure with this spot and its direct neighbours.

For every fitting step, only the current spot with its direct neighbours is fitted. The indirect neighbours are instead added to the model with initialised parameters that are not fitted. Indirect neighbours’ parameter initialisation is performed with an initial guess or with parameters from the previous iteration (when the indirect neighbour was a spot being fitted). After fitting the current spot, SpotMAX moves to the next spot in the sorted list until there are no more spots to fit.

### Benchmark

#### Ground truth 3D dataset

The manually annotated 3D dataset is publicly available at the following link https://hmgubox2.helmholtz-muenchen.de/index.php/s/BFpbnnNGNFQCTk3 and it is composed of three main categories: 1) single-molecule FISH of two genes in *S. cerevisiae*, 2) mitochondrial DNA and mitochondrial network in *S. cerevisiae*, and 3) chromosomal crossovers (tagged COSA-1) in *C. elegans*. The folder structure is compatible with the Cell-ACDC software^9^ and organised into Position folders. The images are in TIFF format, one per channel. Each dataset has been annotated by an expert in each field using Cell-ACDC. The annotations are available in the CSV file ending with “_gt_manual.csv”, while the segmentation masks of the cells are available in the file ending with “_segm.npz”. Additional details about the dataset are available in Supplementary Table 1.

#### Methodology

To match predicted spot coordinates with the ground truth annotations, we paired each predicted spot centre with the ground truth if they were both inside a spheroid with (3, 4, 4) pixels radii in the (z, y, x) directions centred on the ground truth spot. If multiple predicted spots were found in the spheroid, only the closest one was paired. The number of paired spots is the number of true positives (TP), unpaired predicted spots are the false positives (FP), and unpaired ground truth spots are the false negatives (FN). We then calculated precision (P), recall (R), and F1-score (**Fig. 2**) with the following formulas:

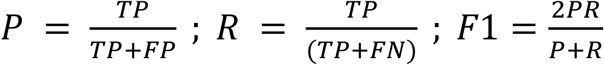

For Spotiflow we used the pre-trained general model^2^, while the RS-FISH analysis parameters are reported in Supplementary Table 5. The SpotMAX analysis parameters for each dataset are available in the published dataset.

### COSA-1 experiments in *C. elegans*

#### Strain

The *C. elegans* strains used for this analysis are the AV630, SMN333, SMN260, SMN350, SMN384, and SMN387. The full genotype of the strains is available in Supplementary Table 4. All strains were maintained at 20 °C on nematode growth medium plates with *E. coli* OP50 as previously described^46^. New strains for this study were generated by CRISPR/Cas9 mediated genome editing as previously described^47^. Sequences are listed in Supplementary Table 6. Cas9-NLS protein, tracrRNA, gRNAs and gBlocks to generate repair templates were purchased from IDT, all oligos were purchased from Sigma Aldrich. To create the *v5::rad-51(ske48)* allele we replaced an HA-tag of a *ha::rad-51(ske1)* strain that we generated using a modified Cas9 to recognise a GAG-PAM motif in the N-terminus of *rad-51* (Supplementary Table 6).

#### Cytology and fixed-tissue imaging

Young adult animals were dissected 24 hours after reaching the L4 stage, fixed and stained (GFP::COSA-1 or V5::RAD-51) as previously described^48^, and mounted in ProLong Glass antifade mounting medium (Invitrogen, P36984). Primary antibodies used for staining GFP and V5 are mouse-anti-GFP (Roche, #11814460001) and mouse-anti-V5 (Invitrogen, #R960-25), respectively, and the secondary antibody is donkey-anti-mouse Alexa Fluor™ 546 (Invitrogen, #A10036). Halo::COSA-1 was labelled with Janelia Fluor 669 HaloTag ligand (Lavis Lab, JBG-27-075A)^49^ by feeding as previously described^50^.

Samples were imaged on an Olympus spinning disk confocal system IXplore SpinSR using a 60x 1.4 NA oil plan-apochromat objective with the cellSens software (Olympus/Evident).

#### Cell segmentation

Meiotic nuclei were segmented as previously described^26^ using a custom model available at https://github.com/KoehlerLab/Cellpose_germlineNuclei/blob/main/Cellpose_germlineNuclei/cellpose_germlineNuclei_KoehlerLab. Segmented nuclei were manually checked using Fiji and only correctly segmented nuclei in late pachytene, marked by the presence of bright COSA-1 spots, were further analysed by SpotMAX.

### *S. cerevisiae* time-lapse experiments

#### Yeast strain

The yeast strain FPY004 used in this work is based on W303 and was constructed by crossing the parental strains (yCO380, containing LacO arrays in mtDNA^14,30^) and ASY014^14^. The full genotype is listed in Supplementary Table 3.

#### Live-cell 4D fluorescence microscopy

The strain FPY004 was grown at 30 °C in a shaking incubator at 250 rpm (Infors, Ecotron). Cells were pre-cultured in YPD medium for 6 hours, transferred to SCGal medium (synthetic complete medium containing 2% galactose), and grown for at least 36 hours. The OD_600_ was always kept below 0.5 with a final OD of 0.3. For the live-cell microscopy experiment, 1 mL of culture was sonicated for 3 s and 200 μL were loaded into a CellASIC^®^ ONIX2 Y04C microfluidic plate (Merck Millipore) connected to the CellASIC^®^ microfluidic pump (Merck Millipore). After trapping the cells in the microfluidic chamber, SCGal medium was continuously supplied with a constant pressure of 13.8 kPa. Image acquisition was performed with a Zeiss LSM 800 confocal microscope, a 63x/1.4 Oil DIC objective, and the Incubator XLmulti S1, Pecon used to maintain an incubation temperature of 30 °C. Cells were imaged for 3 hours, then the medium was automatically changed to SCD medium (synthetic complete medium containing 2% glucose) and imaged for additional 4.5 hours. The acquisition rate was set to 15 minutes. Multi-dimensional imaging was achieved by acquiring 24 z-slices with 0.35 μm spacing for the 3 channels named T-PMT, mNeonGreen, and mKate2. mKate2 was imaged with an excitation wavelength of 561 nm, and detected emission between 610 and 700 nm. mNeon was excited at 488 nm and detected between 410 and 546 nm. Bright-field images were taken using the transmitted light detector (T-PMT). All channels were acquired at the same time.

#### Cell segmentation, tracking, and cell cycle annotation

All the image analysis steps before SpotMAX analysis were performed with the software Cell-ACDC^9^. Raw microscopy files generated by the Zeiss microscope were converted into TIFF format using the BioFormats library^51^. Cells were segmented and tracked using YeaZ^52^ v1.0.3 (embedded in Cell-ACDC) from the bright-field channel. Each time point was then carefully inspected and corrected. The cell cycle annotation was then performed by annotating mother-bud pairing, bud emergence, and cell division. This information was then fed into SpotMAX to count the number of spots (nucleoids) and segment the mitochondrial network in each cell.

### Telomere and centromere fluorescent *in situ* hybridization (FISH) on HSCs

All work was performed in accordance with the Massachusetts Institute of Technology (MIT) Institutional Animal Care Facility and with guidelines at MIT (Institutional Animal Care and Use Committee) (protocol numbers 0715-073-18 and 0718-053-21). Mice were purchased from Jax Laboratories C57BL/6 J (#000664).

Murine bone marrow (BM)-derived live G0/1 HSCs (Lin^-^, Sca1/Ly6^+^, CD117/cKit^+^, CD150/Slamf1^+^, CD48/Slamf2^-^, 7-ADD^-^) were isolated as described previously^30^. Briefly, BM was harvested by flushing the long bones. Red blood cells were lysed in ammonium-chloride-potassium (ACK) buffer and samples were washed in Iscove’s modified Dulbecco’s medium (IMDM) containing 2% foetal bovine serum (FBS). BM cells were resuspended at 10 cells/mL in pre-warmed IMEM supplemented with 2% FBS and 6.6 μg/mL Hoechst-33342 (Thermo Fisher Scientific, #H3570). After 45 min of incubation at 37 °C in a water bath, cells were washed with cold IMEM with 2% FBS and kept at 4 °C. Lineage-positive cells were depleted using a mouse lineage cell depletion kit and the following antibodies were used for staining: Rat monoclonal PE anti-mouse CD150, BD Biosciences, Cat#562651; RRID: AB_2737705; Rat monoclonal APC anti-mouse CD117, BD Biosciences, Cat#561074; RRID: AB_10563203, Armenian hamster monoclonal APC/Cy7 anti-mouse CD48, BioLegend, Cat#103431; RRID: AB_2561462, Rat monoclonal BV711 anti-mouse Ly-6A/E, BioLegend, Cat#108131; RRID: AB_2562241. Cells were sorted using an Aria cell sorter (Becton Dickinson).

HSCs were resuspended in PBS and mounted to a well of a 4-well chamber slide (MatTek) coated with poly-L-lysine for 1 hour at room temperature. HSCs were then washed with PBS twice for 2 minutes each time on a slow rocking platform. HSCs were treated with freshly made pepsin solution (0.005% pepsin in 10 mM HCl) before the FISH procedure. The telomere probe (TelC-Alexa488 [target sequence: telomeric CCCTAA repeats]) and centromere probe (CENPB-Cy3 [target sequence: ATTCGTTGGAAACGGGA]) were mixed and diluted to 500 nM per probe, in a hybridization buffer (20 mM Tris, pH 7.4, 60% formamide, 0.5% of Roche blocking reagent [Roche 11096176001]) for hybridization. 400 µL of diluted probe was used per well (per sample). The rest of the FISH procedure was performed according to the PNA FISH manufacture protocol (PNA Bio). The slides were air-dried briefly before being applied with the Prolong Gold Antifade Mountant medium (Thermo Fisher Scientific) and covered with a 1.5 coverslip. The slides were sealed with nail polish and kept in the dark overnight before imaging. An Ultra Deltavision fluorescent microscope with a 60x objective lens (1.4 NA) was used to acquire the images of nuclei with FISH-labelled telomeres and centromeres, under the setting of 1X1 binning and a z-stack of 20 slices with 0.75 µm spacing size. During the analysis, we excluded cells that were larger than 2.5 SD than the mean cell size. To analyse HSCs of a specific size, we evaluated the 10% smallest (XS-HSCs), the 10% largest (XL-HSCs) and +/− 10% HSCs of mean size (M-HSCs).

### Nucleolus and nuclear size analysis in *C. elegans*

#### Strain

The strain DCW279 was used for this analysis. The full genotype is available in Supplementary Table 4.

#### Auxin plates and treatment

Auxin 3-Indoleacetic acid (IAA) (Sigma-Aldrich) was dissolved in ethanol to prepare a 57 mM stock solution and stored at 4 °C. Auxin was added to NGM plates to a final concentration of 1 mM. Control plates contained an equivalent amount of ethanol (1.75%). Plates were then seeded with OP50 bacteria and let dry.

Embryos isolated by standard hypochlorite treatment were plated onto auxin, or ethanol plates, as control, and maintained for 24 hours. Next, the hatched L1 larvae were collected from the plates and imaged.

#### Microscopy

Microscopy was performed using a confocal spinning disk microscope system from Visitron Systems GmbH, equipped with a Nikon Eclipse Ti2 microscope, aPlan Apo λ 100x/1.45 oil objective, a Yokogawa CSU-W1 confocal scanner unit, a VS-Homogenizer, an EMCCD camera [Andor - iXon Series], and VisiView software for acquisition.

Live microscopy was carried out on 2% agarose pads supplemented with 0.15 % sodium azide (Interchim) to paralyse the worms, as previously described^41^. For each image, a range of 70-75 stacks were captured with a z-spacing of 200 nm.

## Acknowledgements

We thank the Advanced Light Microscopy Facility (ALMF) at the European Molecular Biology Laboratory (EMBL) in Heidelberg, Evident/Olympus and Zeiss for their support; IT and HPC resources at the European Molecular Biology Laboratory (EMBL) in Heidelberg for providing the essential computational infrastructure; the Media and Lab Kitchen at the European Molecular Biology Laboratory (EMBL) in Heidelberg for providing media and solutions. Some strains were provided by the CGC, which is funded by the NIH Office of Research Infrastructure Programs (P40 OD010440). We also thank Björn Schumacher (University of Cologne) for providing essential *C. elegans* strains for this study. Work in the K.M.S laboratory was supported by the Human Frontier Science Program (career development award to K.M.S), the Deutsche Forschungsgemeinschaft (DFG, German Research foundation) through projects 416098229 and 431480687, by the Helmholtz Association, and by the de.NBI Cloud within the German Network for Bioinformatics Infrastructure (de.NBI) and ELIXIR-DE (Forschungszentrum Jülich and W-de.NBI-001, W-de.NBI-004, W-de.NBI-008, W-de.NBI-010, W-de.NBI-013, W-de.NBI-014, W-de.NBI-016, W-de.NBI-022). Work in the J.L. lab was supported by the European Research Council, the Research Council of Finland and the Swedish Research Council. Work in the X.A.S. lab was supported by a Jane Coffin Childs Memorial Fellowship, the Centre for Prostate Disease Research core funding at the Henry Jackson Foundation, and the Genitourinary Malignancies Branch of the National Cancer Institute at the National Institutes of Health. Work in the S.K. lab was supported by the European Molecular Biology Laboratory, and by the Deutsche Forschungsgemeinschaft (DFG, German Research Foundation) - project number 452616889.

