## Supplemental Information for "SpotMAX: a generalist framework for multi-dimensional automatic spot detection and quantification"

### Supplementary Information

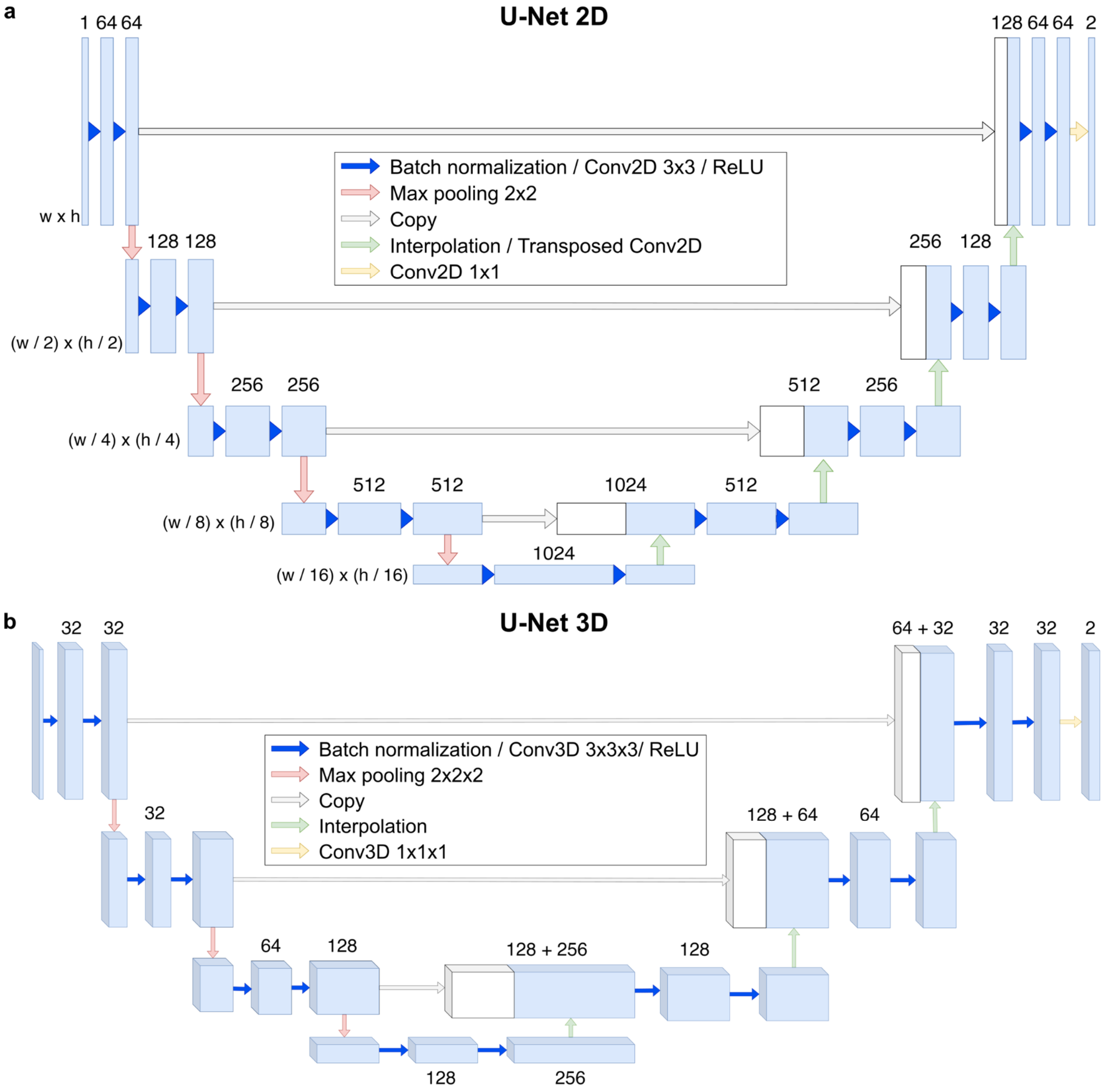


**Supplementary Fig. S1 | Model architectures of the SpotMAX U-Net models**. Blocks represent feature maps, while the number on top is the number of filters. **a**, U-Net 2D. Compared to the original U-Net, we use padding during convolution to avoid decreasing the image size. **b**, U-Net 3D based on the library pytorch3d-unet.

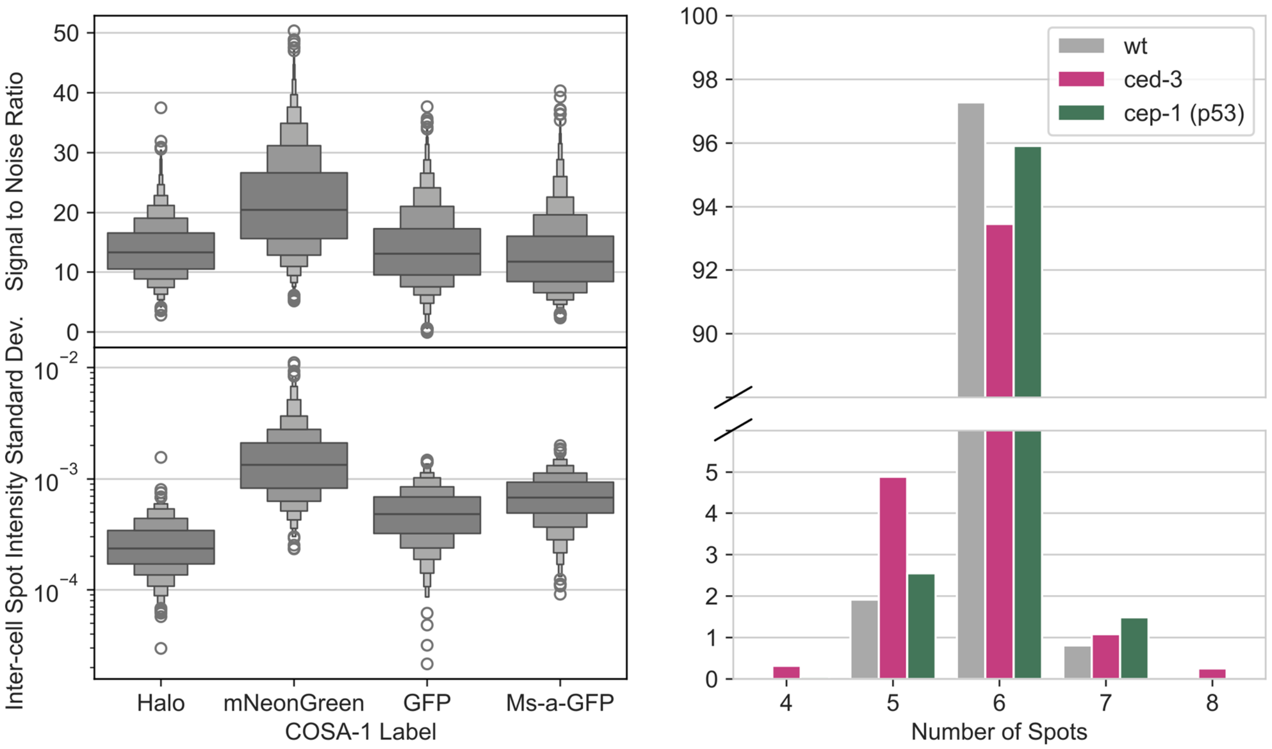


**Supplementary Fig. S2 | SpotMAX analysis of COSA-1 labels and crossover counting in *C. elegans* apoptosis mutants**. **a**, Signal-to-Noise Ratio (SNR) of each spot (top) and standard deviation of the spot intensity distribution in each nucleus (bottom) for the 4 visualisation methods tested (see main text). SpotMAX calculates the SNR as the difference between the mean of the spot intensities within the resolution-limited volume and the mean of the background all divided by the standard deviation of the background (i.e. Glass's effect size). The background was defined as those pixels outside the spot masks and inside the nuclear masks. **b**, Number of chromosomal crossovers in all the segmented nuclei without excluding nuclei positive for RAD-51 (i.e., without unresolved DNA breaks).

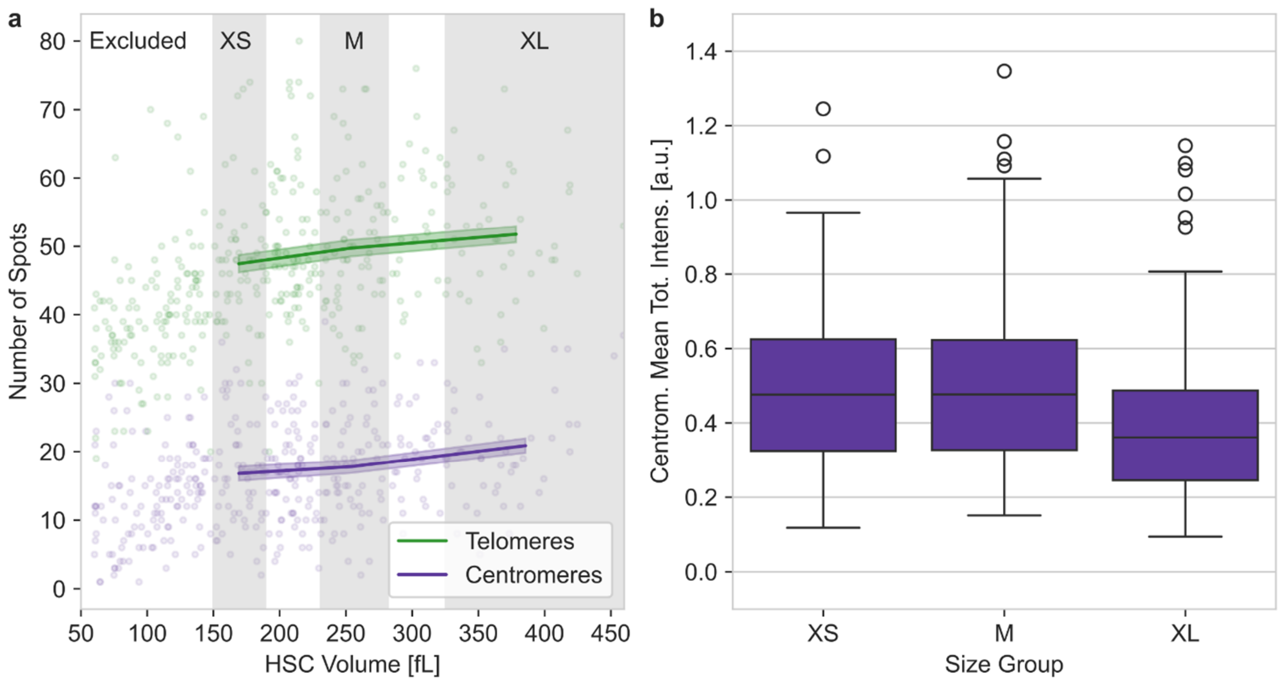


**Supplementary Fig. S3 | SpotMAX analysis of telomeres and centromeres from DNA-FISH data in hematopoietic stem cells**. **a**, Number of spots for both telomeres and centromeres as a function of the entire range of analysed hematopoietic stem cells. In the main analysis, cells below 150 fL were excluded because the number of spots was increasing with cell volume, indicating a higher degree of telomeres and centromeres clustering in the smaller cells. **b**, Centromeres mean total intensity in each cell for each size category group. Number of cells per category: Excluded = 123, XS = 55, M = 65, XL = 55, Total = 175.

| Dataset | Organism | Method | Number of volumes | Number of cells | Gene | Pixel size (pixel/µm) | z-space (µm) | Figure |
|---|---|---|---|---|---|---|---|---|
| Yeast_ smFISH | *S. cerevisiae* | Single-molecule RNA FISH | 51 | 373 | *MDN1* | 0.0721 | 0.24 | 2a,b |
| | | | 109 | 1616 | *SUT509* | | | |
| Yeast_ mito | *S. cerevisiae* | LacO-LacI-mNeonGreen | 190 | 357 | mtDNA (LacO) | 0.0672 | 0.35 | 2c,d |
| C_elega ns_CO | *C. elegans* | (Endogenous) tagging or antibody staining | 19 | 548 | *cosa-1* | 0.0339 | 0.15 | 2e,f |

**Supplementary Table 1 | Experimental datasets used to benchmark SpotMAX**. “mtDNA” stands for mitochondrial DNA, and LacO are the Lactose Operon repeats stably integrated into mtDNA. More details about the mtDNA visualisation system are available in Ref. [14,30].

| Dataset | Organism | Method | Number of volumes | Number of cells | Gene | Pixel size (pixel/µm) | z-space (µm) | Figure |
|---|---|---|---|---|---|---|---|---|
| C_elegans_ CO_labels | *C. elegans* | (Endogeno us) tagging or antibody staining | 58 | 1687 | *cosa-1* | 0.0339 | 0.15 | 3b,c,d |
| C_elegans_ CO_apopt | *C. elegans* | Endogeno us tagging | 120 | 3399 | *cosa-1* | 0.0339 | 0.15 | 3e,f,g |
| Yeast_mito _timelapse | *S. cerevisiae* | LacO-LacI-mNeonGre en | 223 | 280 | mtDNA (LacO) | 0.0792 | 0.35 | 4 |
| HSCs_telo_ centromere | Murine Hematopo ietic stem cells | DNA FISH | 292 | 298 | Telomer es and centrome res repeat | 0.0547 | 0.75 | 5 |
| C_elegans_ Raptor | *C. elegans* | Endogeno us tagging | 41 | 715 | *fib-1* | 0.1346 | 0.2 | 6 |

**Supplementary Table 2 | Experimental datasets used to test SpotMAX's ability to obtain new biological insights**. “mtDNA” stands for mitochondrial DNA, and LacO are the Lactose Operon repeats stably integrated into mtDNA. More details about the mtDNA visualisation system are available in Ref. [14,30].

| Name | Genotype | Description | Origin | Figure |
|---|---|---|---|---|
| FPY004-1 | ***Mat α/a;*** *ura3/URA3, TRP1/TRP1, mt-LacO, ADE2/ADE2, HO/HO::CUP1prom-SU9-2xmNeonGreen-LacI-PGK1prom-SU9-mKate2-KanMX4* | Diploid microscopy strain with LacO-LacI system | This study | 4 |

**Supplementary Table 3 | Yeast strain used in this work**. All strains are based on W303.

| Name | Strain | Genotype | N (animals) | Viability | p-value | Incidence of males [%] | p-value | Origin | Figure |
|---|---|---|---|---|---|---|---|---|---|
| N2 | N2 | N2 | 2660 (10) | 105 ± 3.8 | | 0.173 ± 0.185 | | | |
| *GFP::cosa-1* | AV630 | *meIs8 [pie-1p::GFP::cosa-1 + unc-119(+)] II* | | n.d. | | n.d. | | Yokoo R. et al[17] | 3 |
| *cosa-1::mNeonGreen* | SMN333 | *cosa-1::mNeonGreen(ske21.1) III* | 1322 (6) | 112 ± 5.2 | 0.531 | 0.055 ± 0.136 | 0.245 | This study | 3 |
| *Halo::cosa-1* | SMN260 | *Halo::cosa-1(ske25) III* | 1548 (6) | 104 ± 5.8 | 0.931 | 0.457 ± 0.28 | 0.074 | This study and Neves A.R.R. et al[50] | 2e,f<br>3 |
| wild-type | SMN350 | *Halo::cosa-1(ske25) III; V5::rad-51(ske48) IV* | 1181 (6) | 103 ± 8 | 0.806 | 0 ± 0 | 0.074 | This study | 3 |
| *cep-1* | SMN384 | *cep-1(lg12501) I; Halo::cosa-1(ske25) III; V5::rad-51(ske48) IV* | | n.d. | | n.d. | | This study | 3 |
| *ced-3* | SMN387 | *Halo::cosa-1(ske25) III; ced-3(ske28) V5::rad-51(ske48) IV* | | n.d. | | n.d. | | This study | 3 |
| *daf-15::AID* | DCW279 | *daf-15(re257[daf-15::mNG::AID]) IV; wrdSi23 [eft-3p::TIR1::F2A::BFP::AID*::NLS ::tbb-2 3'UTR] I; mcherry::fib-1 (ustIS36) II; his-72 (uge30[gfp::his-72]) III* | | n.d. | | n.d. | | This study | 6 |

 **Supplementary Table 4 | *C. elegans* strains used in this study**

| | **Yeast_smFISH -** *MDN1* | **Yeast_smFISH -** *SUT509* |
|---|---|---|
| **Parameter name** | **Value** | **Value** |
| SigmaDoG | 1.5 | 1.5 |
| ThresholdDoG | 0.0025557573 | 0.007831375 |
| anisotropyCoefficient | 4.02 | 1.3730820417404175 |
| useAnisotropyForDoG | true | true |
| RANSAC | SIMPLE | SIMPLE |
| MaxError | 1.5 | 1.5 |
| InlierRatio | 0.108 | 0.1 |
| supportRadius | 3 | 3 |
| Intensity computation | Gaussian fit (on inlier pixels) | Linear Interpolation |
| min intensity | 816.0 | 838.0 |
| max intensity | 2683.0 | 7522.0 |
| autoMinMax | true | true |
| bsMethod | RANSAC on Mean | No background subtraction |
| bsMaxError | 0.05 | 0.05 |
| bsInlierRatio | 0.1 | 0.1 |

**Supplementary Table 5 | RS-FISH analysis parameters used for the benchmark in Fig. 2 in the main text**.

| Allele | crRNA sequence | Repair template sequence | Genotyping oligos |
| --- | --- | --- | --- |
| *cosa-1::mNeonGreen(ske21.1) III* | tgtcagagatggtagTTACG | GgtaactgctggccgagtttttctaggccacgcgtggcaattttacaattaattatttttatttatttcagAATGAGAGTATTCCGGAATGCAGCACCTCCTCGGTCTCCAAGGGAGAGGAGGACAACATGGCCTCCCTTCCAGCCACCCACGAGCTTCACATCTTCGGATCCATCAACGGAGTCGACTTCGACATGGTCGGACAGGGAACCGGAAACCCAAACGACGGATACGAGGAGCTTAACCTTAAGTCCACCAAGgtaagtttatctaaaaagtttttcattcaaaatgtgtaaaaattcatttaaaataacccaaaaatcattaatcctcgatattttcagGGAGACCTTCAATTCTCCCCATGGATCCTTGTCCCACACATCGGATACGGATTCCACCAGTACCTTCCATACCCAGACGGAATGTCCCCATTCCAAGCCGCCATGGTCGACGGATCCGGATACCAAGTCCACCGCACCATGCAATTCGAGGACGGAGCCTCCCTTACCGTCAACTACCGCTACACCTACGAGGGATCCCACATCAAGGGAGAGGCCCAAGTCAAGGGAACCGGATTCCCAGCCGACGGACCAGTCATGACCAACTCCCTTACCGCCGCCGACTGGTGCCGCTCCAAGAAGACCTACCCAAACGACAAGACCATCATCTCCACCTTCAAGTGGTCCTACACCACCGGAAACGGAAAGCGCTACCGCTCCACCGCCCGCACCACCTACACCTTCGCCAAGCCAATGGCCGCCAACTACCTTAAGAACCAGCCAATGTACGTCTTCCGCAAGACCGAGCTTAAGCACTCCAAGACCGAGCTTAACTTCAAGGAGTGGCAGAAGGCCTTCACCGACGTCATGGGAATGGACGAGCTTTACAAGggaggtggcTAActaccatctctgacagcacctctttgtcgccgattccactggtcgcggctcgttcactgcaacaaattattgatttttattgtcatgtaccatattg | FW:<br>GCAATCGATTATAGAGCTCGCC<br>RV:<br>tgcaatgagtacgtgacagg |
| *ha::rad-51(ske1) IV* | TTTCTTTTGACGACTTGCT<br>(GAG crRNA) | ccaattgtatgttttaacttaaaaataattaattaaaattatctcaggagtcaaaaATGTACCCCTACGATGTCCCAGATTATGCTTCAGCaCAAGCAAGTCGTCAAAAGAAATCGGATCAAGAGCAGCGTGC | |
| *V5::rad-51(ske48) IV* | CATAATCTGGGACATCGTAG<br>(targeting HA-tag) | cttaaaaataattaattaaaattatctcaggagtcaaaaATGGGAAAGCCAATCCCAAACCCACTTCTTGGACTCGACTCCACCTCAGCaCAAGCAAGTCGTCAAAAGAAATCGGATCAA | FW:<br>gctccactgttaaaaatgccg<br>RV:<br>ATATCGCCAGAACTGATTCCTG |
| *ced-3(ske28)* | ataaatttttgcagcaagtg | CatttttaaatgataattaataaatttttgcaaCAAGTGatcAGAAAGAAGCCGAGCCAAGCTg | FW: ggtcgcagttttcagtttagag<br>RV:<br>CCACAAGCGACCTTCTTATTG |

**Supplementary Table 6 | List of sequences.** Sequences used to generate *cosa-1::mNeonGreen* and *Halo::cosa-1* strains (Supplementary Table 4) with CRISPR/Cas9.
